# Intestinal flow rates, absorption of felodipine from the small intestine and attributes of chyme collected at midgut from Labradors

**DOI:** 10.1101/2024.01.05.574377

**Authors:** Steffen M. Diebold, In memory of Mia Wilsson-Rahmberg

## Abstract

The objectives of the present study were (1) to investigate gastrointestinal hydrodynamics of Labradors as a model for human midgut (2) to examine various attributes of intestinal fluids in vivo and (3) to study the influence of hydrodynamics on the dissolution and absorption of a poorly soluble drug from various suspensions.

Gastrointestinal flow rates were determined volumetrically using an aspiration method. Isotonic saline and 20 % glucose solutions were used to alter gastrointestinal hydrodynamics. Felodipine, a BCS class II substance, was suspended in these fluids. Osmolality, pH, bile acid concentration and drug solubility in various chyme samples were determined. Blood plasma levels of felodipine were recorded while gastrointestinal dissolution was ongoing.

Fluid recovery at midgut fistula was significantly higher (>100 %) for glucose 20 % than for isotonic saline solutions (70 %). After administration of 200 ml glucose 20 % the (overall) grand median of differential gastrointestinal flow rates (DFR) was 8.3 ml/min.. Individual spike flow ranged from 20 up to 60 ml/min. Corresponding flow rates after administration of 200 ml isotonic saline were 35.0 ml/min. for the grand median including individual spike flows beyond 100 ml/min.. Within and between-dog variability in flow rate data was similar. In general, glucose solutions released more evenly. Following oral administration of glucose solution 20 % osmolality of intestinal fluids decreased within 40 min. from about 1000 mOsm. towards more physiological values of about 350 mOsm.. Saturation solubility of felodipine (C_s_) in jejunal chyme after administration of either solution (saline or glucose) was determined to be about 10 (µg/ml) on average (median), *exposing high variability with time*! The intestinal solubility varied greatly within the course of an experiment. However, a strong correlation was observed between the aspirated fluid volume and the dissolved amount of felodipine confirming the well known relationship of Noyes, Whitney, Nernst and Brunner in-vivo. Grand median of pH in jejunal chyme of labradors was determined to be 6.68. Median values range from 4.38-7.62. The pharmacokinetic data showed a slight trend to differences based on particle size and on fluid administered.

## 1 Introduction

Many orally administered drugs are poorly soluble but well absorbable in vivo. A variety of factors influence intestinal dissolution and subsequently absorption. Among these are found the gastrointestinal motility, affecting intestinal flow rates, as well as the composition of the gastrointestinal fluids. Since only a few data exist for Labradors on this matter so far, the present study focuses the investigation of intestinal fluids in the upper gastrointestinal tract with respect to flow rates, osmolality, pH, bile salt concentration and drug solubility.

## 2 Materials and Methods

### 2.1 Materials

Felodipine (MW: 384.26 g/mol) was chosen as a model substance for a poorly soluble (aqueous solubility: 1 mg/L at 37°C) but well absorbable drug. Felodipine and the internal standard (H 165/04), a structural analogue of felodipine were supplied by the former Astra Hässle (Mölndal, Sweden). The median particle size of the micronized felodipine powder, Lot# 41688-01, was about 3 µm (d_10_ = 1.38 µm, d_90_ = 9.20 µm). Coarse grade felodipine (Lot# coarseS 200-315) was used with a median particle size of 236 µm (d_10_ = 74 µm, d_90_ = 372 µm). Particle sizes were determined using a Coulter^®^ LS 130 (Coulter Electronics LTD, Florida, USA) equipped with a micro volume module. All other chemicals were of analytical grade or equivalent and purchased commercially.

### 2.2 Animals

Two male Labradors (called “Nubbe” and “Nixon”), aged about more than 1 year and weighing about 29-30 kg were used for the study. The dogs were obtained from Terje, Gammelsrud, Norway. The labradors had a chronic nipple fistula. The outer diameter of the fistula was 2.1 to 2.7 cm. A titanium-V2A-canula was specially handcrafted according to specifications laid down at (Diebold, 2000). The canula (inner diameter 0.9 cm, outer diameter 1.1 cm) was introduced about 5.8 cm into the lumen of the fistula with the help of a protective teflon insert. The experiments were approved by the Ethics Committee of the University of Gothenburg (ethics approval number: 2091997) according to the declaration of Helsinki. The surgery was prepared according to the method of Wilsson-Rahmberg (Wilsson-Rahmberg and Jonsson, 1997).

### 2.3 Study Design

To maintain the fasted state motility pattern a 0.9 % saline solution was given orally via orogastric tubes. To switch to a fed state like motility 20 % glucose solution was administered. Osmolality, ph, bile salt concentration and drug solubility were determined. Intestinal flow rates were measured as well as the absorption of orally administered felodipine suspended in different fluids. In each dog, experiments including felodipine were performed no more than once a week. For pharmacokinetic studies a total of 10.0 mg of felodipine and 200 ml of accompanying fluid were administered.

### 2.4 Preparation of felodipine formulations and administration procedure

#### Solution

10.0 mg felodipine was weighed into a 50 ml Erlenmeyer flask and 2 ml ethanol 99 % (v/v) was added and 20 ml polysorbate 80 (1.5 % (w/v)) to fully dissolve the felodipine. This was followed by 20 ml of a 1.5 % (w/v) HPMC (6cps) matrix solution.

#### Suspension of micronized powder

10.0 mg micronized felodipine was weighed into a 50 ml Erlenmeyer flask and 20 ml of a 1.5 % (w/v) HPMC (6cps) matrix solution was added to prevent any agglomeration. The suspension was sonicated for approximately 1 minute.

About 100 ml of the matrix solutions (0.9 % saline or 20 % glucose solution) were administered via an oro-gastric tube into the stomach. This was followed by 20 ml of the felodipine bolus suspensions (or solution) respectively. Time was started and the remainder of the co-administered matrix solution was given. The administration procedure was completed within 1-2 minutes.

#### Suspension of coarse grade powder

10.0 mg of coarse grade felodipine was weighed in an aluminium boat (1.12 ml) and the boat was carefully transferred into a 50 ml Erlenmeyer flask. The matrix solutions (glucose 20 %, or isotonic saline) were prepared separately. At the time of administration, the orogastric tube was connected to a suitable funnel. The coarse powder was administered via the funnel, then the fluid was used to flush.

### 2.5 Blood samples

Blood samples (2–3 ml) were taken from foreleg veins. Samples were collected at 0, 0.25, 0.5, 1, 1.5, 2, 3, 5, 7 and 24 hours after oral administration into Venoject ꜜ heparin tubes (Terumo, Italy). Samples were stored on ice for no longer than 1-2 hours, then centrifuged (Rotina 48 RS, Hettich, Tuttlingen) for 10 minutes at 4000 rpm and 4°C. The plasma was transferred in a new vial and stored at –20°C until assayed for drug concentration by gas chromatography according to the method of Ahnoff (Ahnoff, 1984). Typical elution times were 13 min and 13.5 min for felodipine and the internal standard, respectively. The dogs had access to food 7 hours after start of the experiments.

### 2.6 Pharmacokinetic Calculations

Pharmacokinetic analysis consisted of the visual identification of the maximum plasma concentration (C_max_) and the time at which this occurred (T_max_) from the individual subject plasma concentration time profiles. The area under the plasma concentration versus time curve from time zero to t (AUC ^t^ ) was calculated using the trapezoidal method. Calculation of the mean values of the parameters and their standard deviation was performed. AUC_24_ und AUC_7_ represent area under time curve from 0 to 24 h (AUC_24_) and 7 h (AUC_7_) after administration of the dose, respectively. Relative bioavailability was calculated according to Frel (%) = (AUC-susp./AUC-oral)*100.

### 2.7 Chyme sampling procedure

The dogs were studied fully conscious while standing or resting comfortably in restraining slings (Wiktor TR 100 - Lyftvagnar, Constella Verken AB, Borlänge, Sweden). Testing always began at about the same time each day. Before each test, the dogs were fasted for 16 hours. Access to water was restricted 1 h before the experiment. Oral intubation was performed with a standard tube (80 cm, Pharma-Plast, Denmark), with the help of a wooden bite-bar to protect the catheter. Syringe and orogastric tube were rinsed twice with the remaining fluid to ensure that all of the substance was transferred into the dog’s stomach. The volume of fluid was administered over a one to two minute period. The orogastric tube was then removed. Samples of chyme were collected with the help of calibrated vials (Venoject, plain not silicone coated, Terumo, Italy) by simple drainage through the fistula until fluid recovery appeared to be complete. Increments of 10 ml of chyme were collected. The vials stick to the titanium canula by means of a plastic ring that is not in contact with the fluid. As none of the dissolved felodipine was ever in contact with materials other than glass or titanium, the likelihood of adsorption was minimized. When introducing the canula into the fistula a teflon inlay with a rounded top was used to protect the mucosa. This inlay was retracted before starting the experiment. The first few ml of chyme that appeared - before administration of the fluid - were discarded. The time to collect each 10 ml increment of fluid was recorded. Centrifugation of chyme (20 C, 5000 g, 8 min) was performed to separate particulate intestinal material and undissolved felodipine from the fluid. The supernatant was transferred into glass vials for later felodipine assay in chyme (=dissolved but not absorbed fraction).

### 2.8 Flow rates and fluid recovery

Calculation of differential flow rates (DFR) in (ml/min.) was performed according to the following equation:

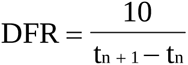

Increments of 10 ml were collected. (t_n+1_ - t_n_) represents the time intervall between two increments. DFRs were averaged over the time period of an experiment to get median differential flow rates. Fluid recovery (V_R_) was referred to the starting volume of 200 ml.

### 2.9 Bile salt analysis

Bile acid assay was performed with a bile salt kit (Enzabile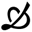, Nycomed, Norway). This test is specific for 3-**✓** -hydroxy-bile acid derivates (BA).

### 2.10 Solubility of felodipine in chyme

Micronized felodipine was added in excess (about 500 to 2500*C_s_) to aliquots of the same samples as used for bile salt determination. The samples were stored and shaken in closed glass vials at 37 °C in an air circulated oven for 3 days until there was no further change in saturation solubility. Prewarmed glass vials, glass syringes (10 mL) and teflon filters (Minisart SRP 25, Sartorius, Göttingen, Germany) were used to separate chyme from precipitated proteins and undissolved felodipine. The filtrate was diluted 1:1 with ethanol 99 % (v/v) to prevent precipitation of felodipine, since the samples had to be stored frozen prior to analysis.

### 2.11 Osmolality

Osmolality of the chyme samples was determined using a calibrated freezing point micro-osmometer (3MO, Advanced Instr. Inc., Mass., USA).

### 2.12 pH-values

The pH-values of chyme samples were determined at room temperature using a calibrated pH-Meter PHM 83 Autocal (Radiometer, Kopenhagen, Danmark).

### 2.13 Statistics

Data are presented as medians and quartiles or, where appropriate, as mean values (±SD). Bifactorial sets of data were compared using bifactorial ANOVA (two prandial states, two dogs). A p-value less than 0.05 was considered significant (SigmaStat 2.0). Appropriate non-parametric tests were used for data that were not normally distributed.

## 3 Results and Discussion

### 3.1 Volume recovery

Volume recovery (V_R_) at midgut after administration of normal saline (200 ml) was generally between 70 and 100 %. After 20 % glucose, more than 100 % of the administered volume was generally recovered, indicating that water efflux occurs into the lumen in response to administration of a hypertonic solution. (For Labrador Nubbe mean values were > 140 %, for Nixon > 150 %). Quantitative fluid recovery is possible using the fistula model. Recovery is in excess of 100 % in the case of 20 % glucose and substantially greater, on average, than for normal saline. Obviously, net water secretion into the lumen of duodenum and jejunum was evoked to compensate for the hypertonic fluids.

### 3.2 Flow rates at midgut

#### 200 ml glucose 20 %

Differential gastrointestinal flow rates after administration of 200 ml glucose solution 20 % generate peak values of 20 up to 60 ml/min. in both labradors (values referred to short periods of time, single experiments). The medians (within single experiments) range from 0.8 ml/min. to 10.9 ml/min. Grand median (all experiments, overall median) was determined to be 8.3 ml/min. The first quartil (Q1) was determined to be 5.2 ml/min. whereas Q3 was 10 ml/min.. In contrast to reported mean transit times ((Diebold, 2000), Chap. VI, 17)) the observed flow rates in this paper were also influenced by the gastric emptying rates.

#### 200 ml isotonic saline

Medians of gastrointestinal flow rates following the administration of normal saline range from 10 to 50 ml/min., the peak maxima from 40 to 200 ml/min. Grand median (all experiments, overall median) was determined to be 35.0 ml/min.. The first quartil (Q1) was determined to be 15 ml/min. whereas Q3 was 40 ml/min.. The observed ranges are relatively narrow for experiments where gastric emptying and midgut transit may both influence the collected volume increments (Diebold, 2000).

#### Variability of differential flow rates

A considerable variability of differential flow rates (DFR) was observed with time (Fig. 1). Variability of DFR occurring in the small intestine of Labradors was more expressed following oral administration of NaCl 0.9 % (Exp.# J) compared to glucose solution 20 % (Exp.# I). This could be attributed in parts to activated gastrointestinal feed back mechanisms after oral administration of caloric fluids and subsequently *a more even release* of the caloric fluid from the stomach.

**Fig 1.**
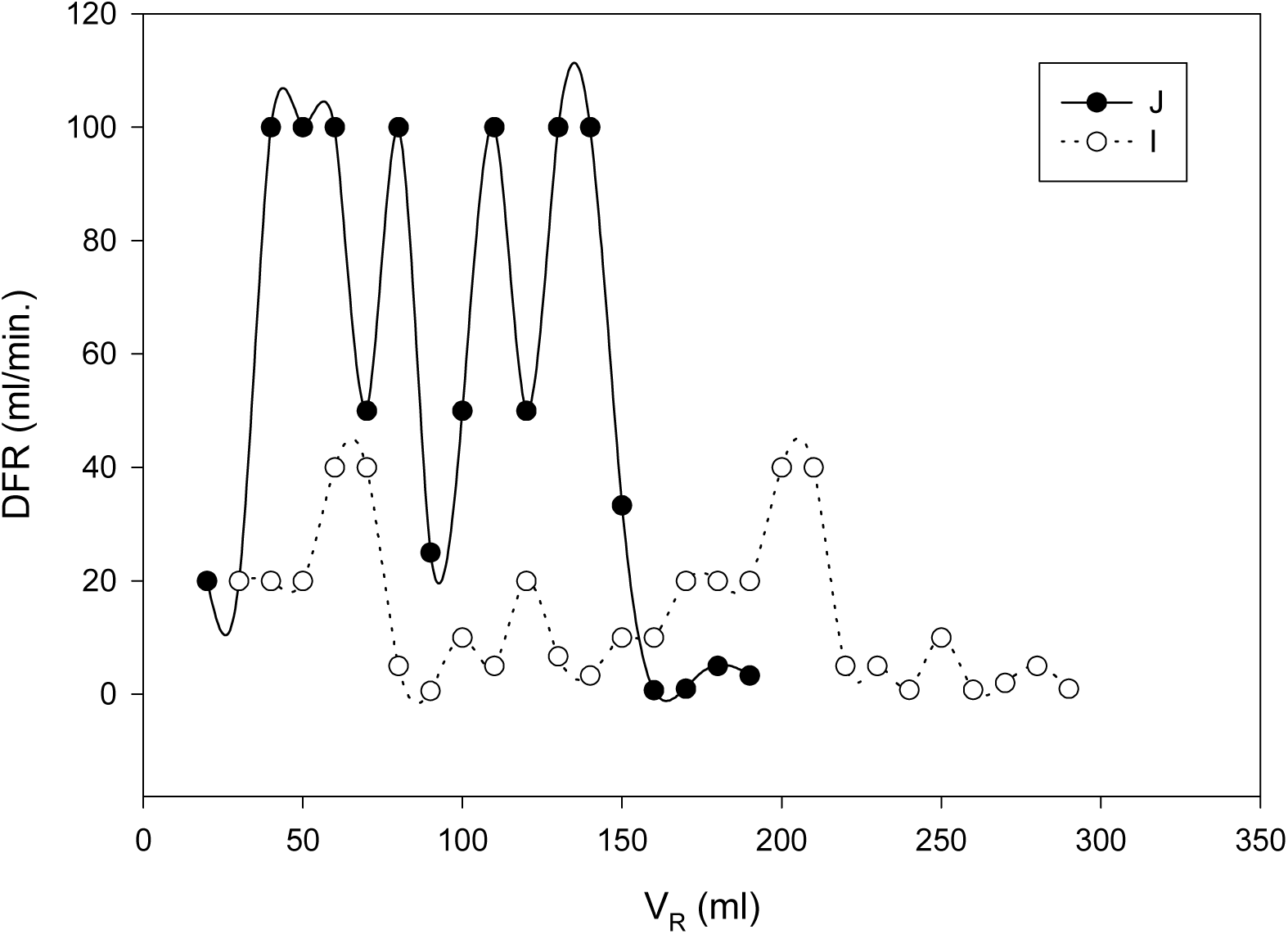
Variability (time dependency) of differential flow rates (DFR) occurring in the small intestine of labradors. (V_R_) represents the volume of chyme collected at midgut (Example given for labrador Nubbe following oral administration of 200 ml glucose solution 20 % (Exp.# I) and 200 ml NaCl 0.9 % (Exp.# J)).

**Fig 2.**
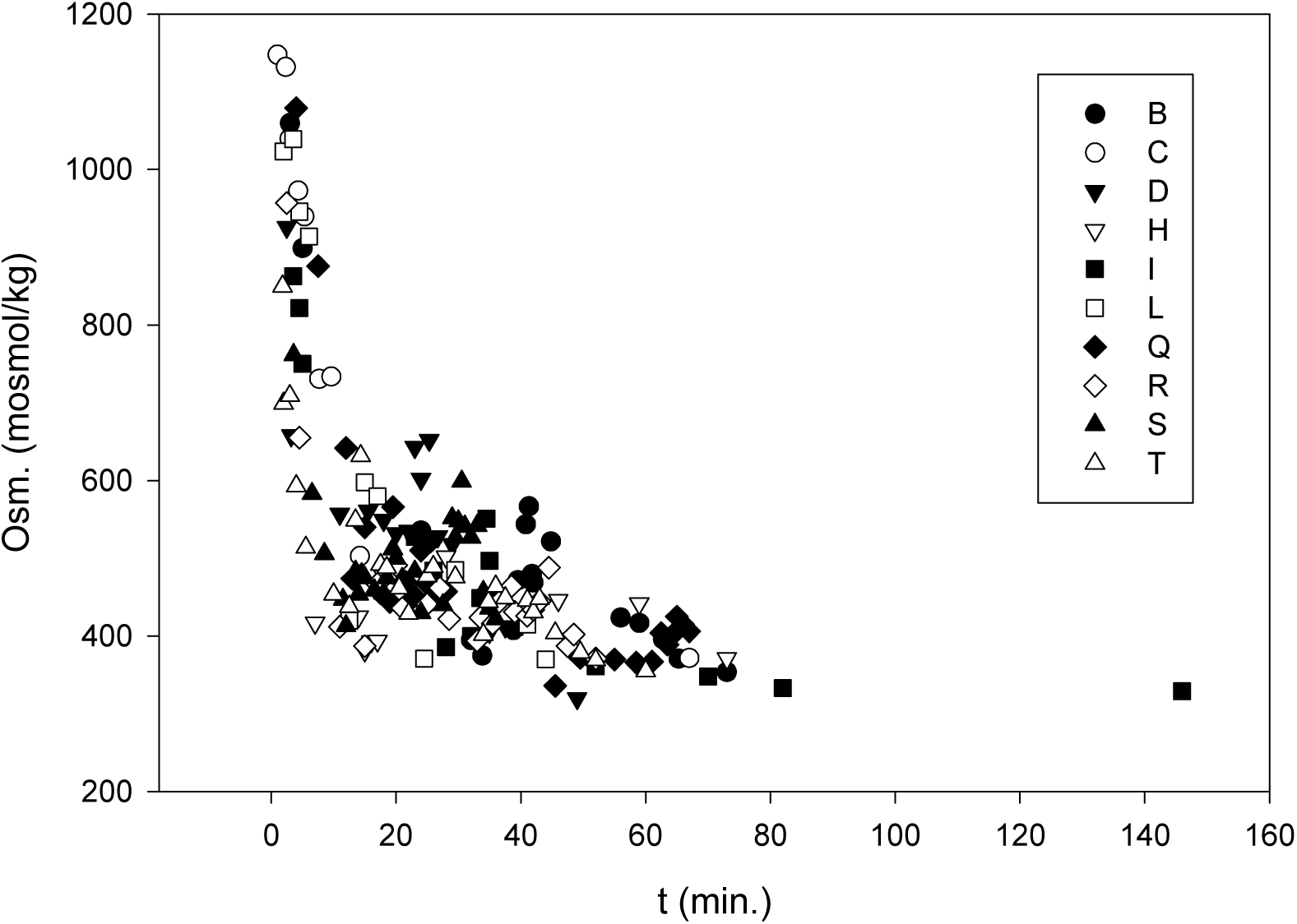
Osmolality of chyme aspirated at midgut for two labradors plotted versus time after oral administration of 200 ml glucose solution 20 %. (190 data from various experiments).

**Fig 3.**
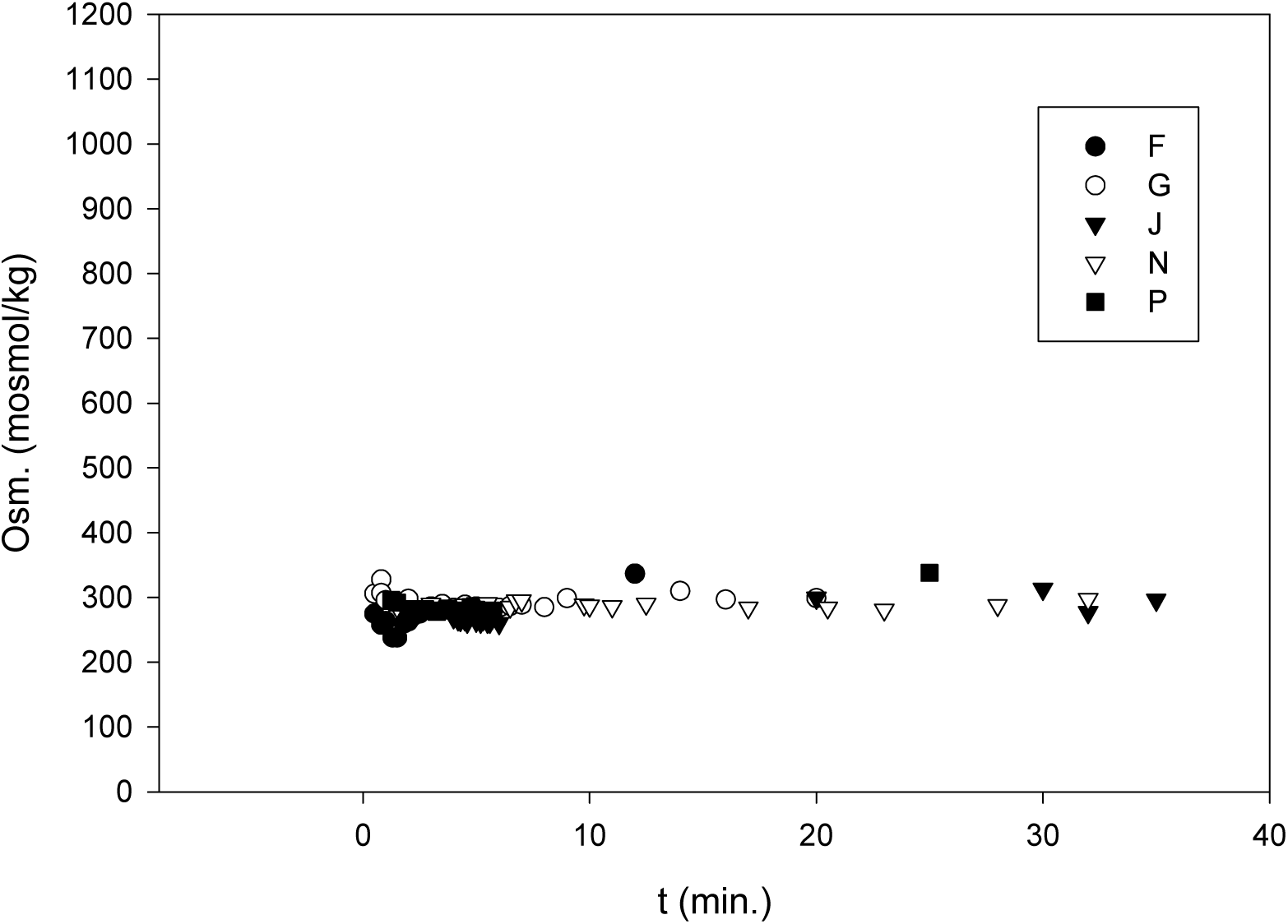
Osmolality of chyme aspirated at midgut. Example given for one labrador (Nubbe) and plotted versus time after oral administration of 200 ml NaCl 0.9 %. (81 data from various experiments).

#### Comparison of treatment

Compared to 200 ml normal saline, the flow rate data for 20 % glucose (200 ml) are considerably slower (8.3 ml/min. vs. 35.0 ml/min., grand medians of differential flow rates). The change that occurred with the treatment is greater than would be expected by chance; there is a statistically significant change (P = 0.018, Mann-Whitney-Test). The difference between the grand median values for 20 % glucose of the two dogs (Nubbe and Nixon) is not great enough to reject the possibility that the difference is due to random sampling variability. There is not a statistically significant difference between the input groups (P = 0.273). In other words, no dog effect was seen in the 20 % glucose data. Compared to intestinal flow rate data from literature for humans (for references see (Diebold, 2000)) our flow rates are faster, probably because of lack of complete feedback effects from ileum and possibly also due to species differences, as well as experimental protocol differences. Conclusion: Flow rate at midgut after administration of 20 % glucose is considerably slower than after normal saline, most probably as a result of decreased gastric emptying (more even release of the caloric fluid from the stomach). 20 % glucose and normal saline can be used to access hydrodynamic effects on dissolution in vivo.

#### Duodeno-jejunal flow rates in the canine small intestine: data from literature

Permanent aspiration of intestinal chyme partially inhibits gastrointestinal feed back mechanisms even in presence of glucose 20 % solutions since ileal glucose receptors are not accessible. This may explain relative high flow rates found in our study for glucose solutions but is no sufficient explanation for surprisingly high flow rates after saline. Nevertheless, other authors reported data that may support our findings and confirm the magnitude of gastrointestinal flow rates in the case of saline solutions: According to data from Greenwood (Greenwood, 1994) mean flow rates of about 40 ml/min. occurred within the first 10 min after oral administration of isotonic saline solutions to dogs (for details on flow rate calculations see (Diebold, 2000)). Sirois (Sirois, 1989) reported median flow rates of 17.3 ml/ min. for the duodenum. However, mean flow rates occurring in the human small intestine appear to be somewhat lower than in the canine small intestine (Oberle et al., 1990).

### 3.3 Osmolality

Nominal osmolality of normal saline was measured to be 283±1 mOsm.. Nominal osmolality of 20 % glucose was measured to be 1172±4 mOsm.. The figures below show osmolality profiles in vivo.

Osmolality data are similar in the two dogs. Our results indicate that osmolality in chyme stays constant at midgut after administration of normal saline (240 to 340 mOsm.). There is a lack of trend with time after administration of normal saline. In contrast, osmolality starts off high (beyond 1000 mOsm.) and gradually trends down to isoosmolar values with time after administration of 20 % glucose. Even 40 min. after administration of 200 ml glucose 20 % osmolality in chyme was measured to be in most cases beyond about 400 mOsm.. The data are fairly consistent with volume/recovery data and suggest that water is transported into the gut lumen to reduce the osmolality when a hyperosmolar solution is administered. (Hendrix et al., *1987;* Reppas et al., 1998), (Trendelenburg, 1917).

### 3.4 pH of intestinal chyme collected at midgut from Labradors

pH values of liquids prior to administration were determined for 0.9 % normal saline to be: 7.42±0.11 (n = 3) and for glucose solution 20 % to be 7.28±0.07 (n = 3). pH-values at midgut after administration of 200 ml of 20 % glucose or normal saline in the two dogs are quite similar. The median values range from 5.93-7.55 for 20 % glucose in the chyme of both Labradors, whereas median values range from 4.38-7.62 for normal saline. There appears to be no difference in pH with fluid administered. There is a lack of trend with time in pH as well. The (overall) grand median for intestinal chyme collected at midgut from Labradors was pH=6.68. For *beagles*, however, many pH-data are found in literature. Youngberg et al. *(Yo*ungberg et al., 1985) performed a radioelemetric investigation (Heidelberg-Kapsel, 7 mm, ante cenam) and found that pH (± SD) was 6.7±0.05 which is comparable to the data reported here for Labradors. Note that because felodipine does not ionize in the physiologically relevant pH range, pH is unlikely to have any effect on either solubility or dissolution for the particular drug used in this study but may be an important factor for ionizable compounds.

### 3.5 Solubility of felodipine in intestinal fluids

The solubility of felodipine was measured in various chyme samples (Fig. 4). In many cases the solubility was much greater than the aqueous solubility (∼ 1 µg/ml). The solubility varied greatly within the course of an experiment (e.g. from 1 µg/ml to 25 µg/ml in one experiment and from 1 µg/ml to 273 µg/ml in another experiment). The cause of increase and variability (time dependency) of the drugs intestinal solubility is at present not clear. The composition of the intestinal fluids may change with time in the course of an experiment. A weak linear correlation could be found for the solubility of felodipine in chyme with bile salt concentration in the case of isotonic saline. However, in the presence of glucose no trend could be found.

**Fig 4.**
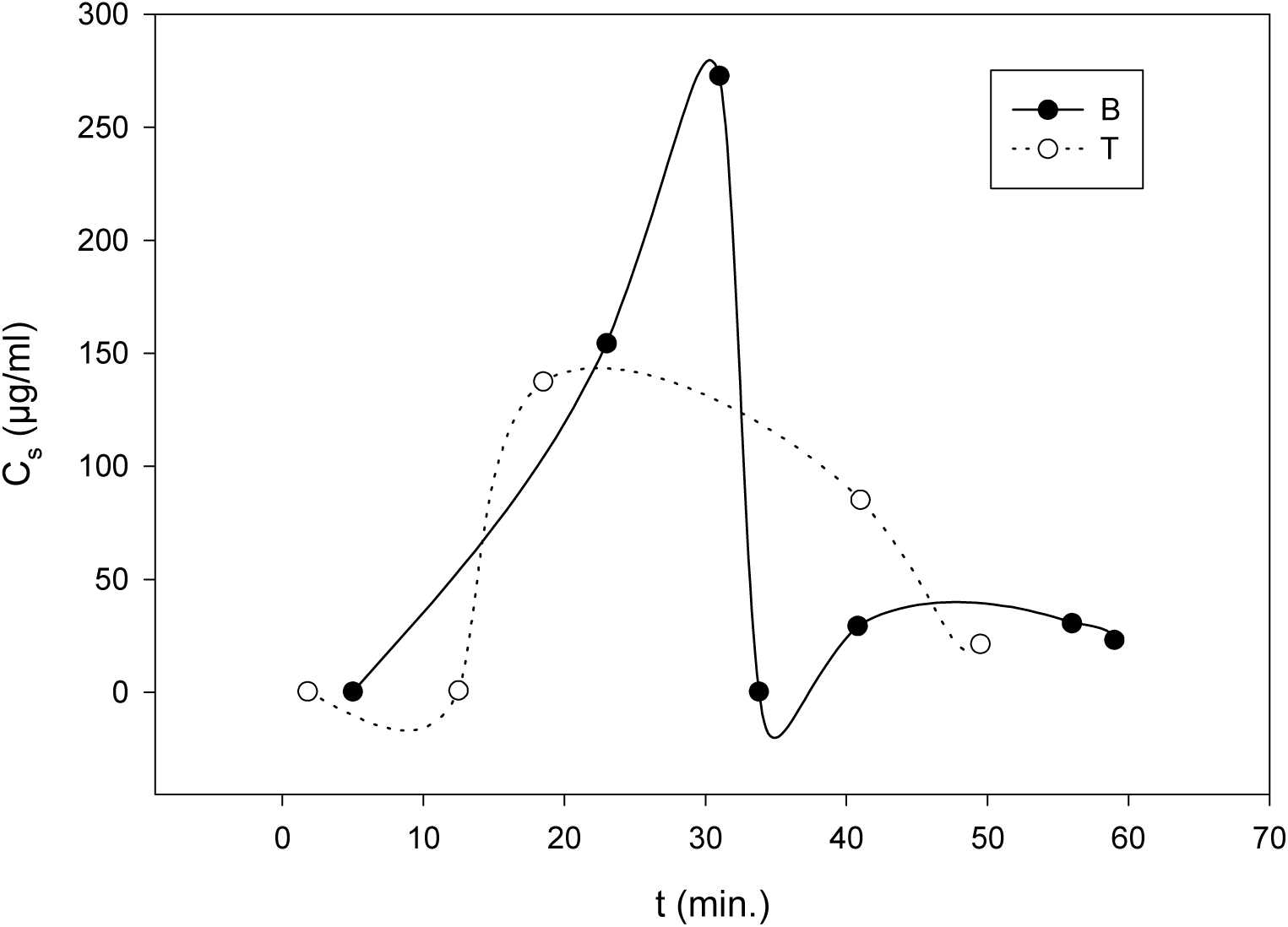
Saturation solubility of felodipine (C_s_) in chyme after administration of 200 ml glucose 20 % solution, depicted for Labradors Nubbe (B) and Nixon (T), showing typical variability with time.

After administration of normal saline grand median for the solubility was determined to be 9.3 (µg/ml), with single experiment medians varying from 0.2 to 42.0 (µg/ml). After administration of glucose solution 20 % grand median was 9.8 (µg/ml) with single experiment medians varying from 0.3 to 29.5 (µg/ml) for Labrador Nubbe and 14.1 (µg/ml) for Nixon, with single experiment medians varying from 0.9 to 112.6 (µg/ml). Maxima of (C_s_) were higher after glucose than after administration of isotonic saline solution. The observed time dependency of intestinal drug solubility casts doubts on most dissolution models that sufficiently describe dissolution rates in vitro but may fail to do so in vivo.

### 3.6 Intestinal bile salt concentration (Bile acid assay)

Difficulties with bile acid assay were encountered, however all results were in the micromolar range, indicating that 200 ml fluid, either saline or 20 % glucose, failed to stimulate bile output in a way similar to a normal meal. This is reasonable since neither liquid contained fats or oils. Therefore, changes in flow rate can be achieved independently of changes in lumenal bile salt concentrations, making it easier to pinpoint hydrodynamic effects on drug dissolution or absorption. In the case of isotonic saline solutions it appears to exist a weak correlation between the solubility of felodipine and the bile salt concentration in chyme (Fig. 5). This trend could not be observed for glucose solutions. Note that secretion of bile acids in dogs is controlled by a circadiane rhythm as well (Kararli, 1995).

**Fig 5.**
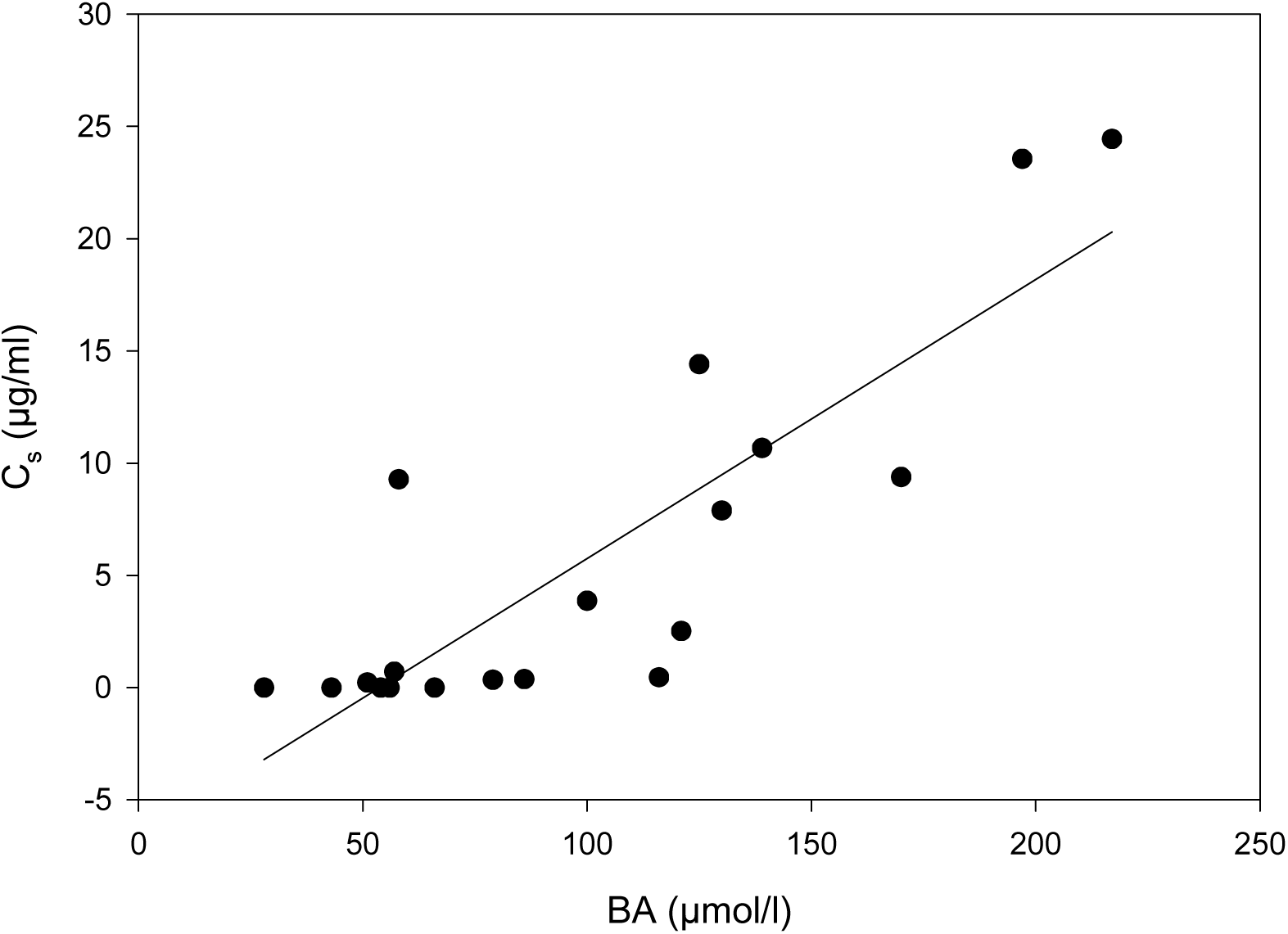
Saturation solubility (C_s_) of felodipine as a function of intestinal bile acid concentration (BA) following the administration of 200 ml NaCL 0.9 % and depicted for labrador Nubbe (C_s_=0.124*BA-6.698, R^2^=0.72). For saturation solubility data of felodipine after administration of glucose solutions see (Diebold 2000).

**Fig 6.**
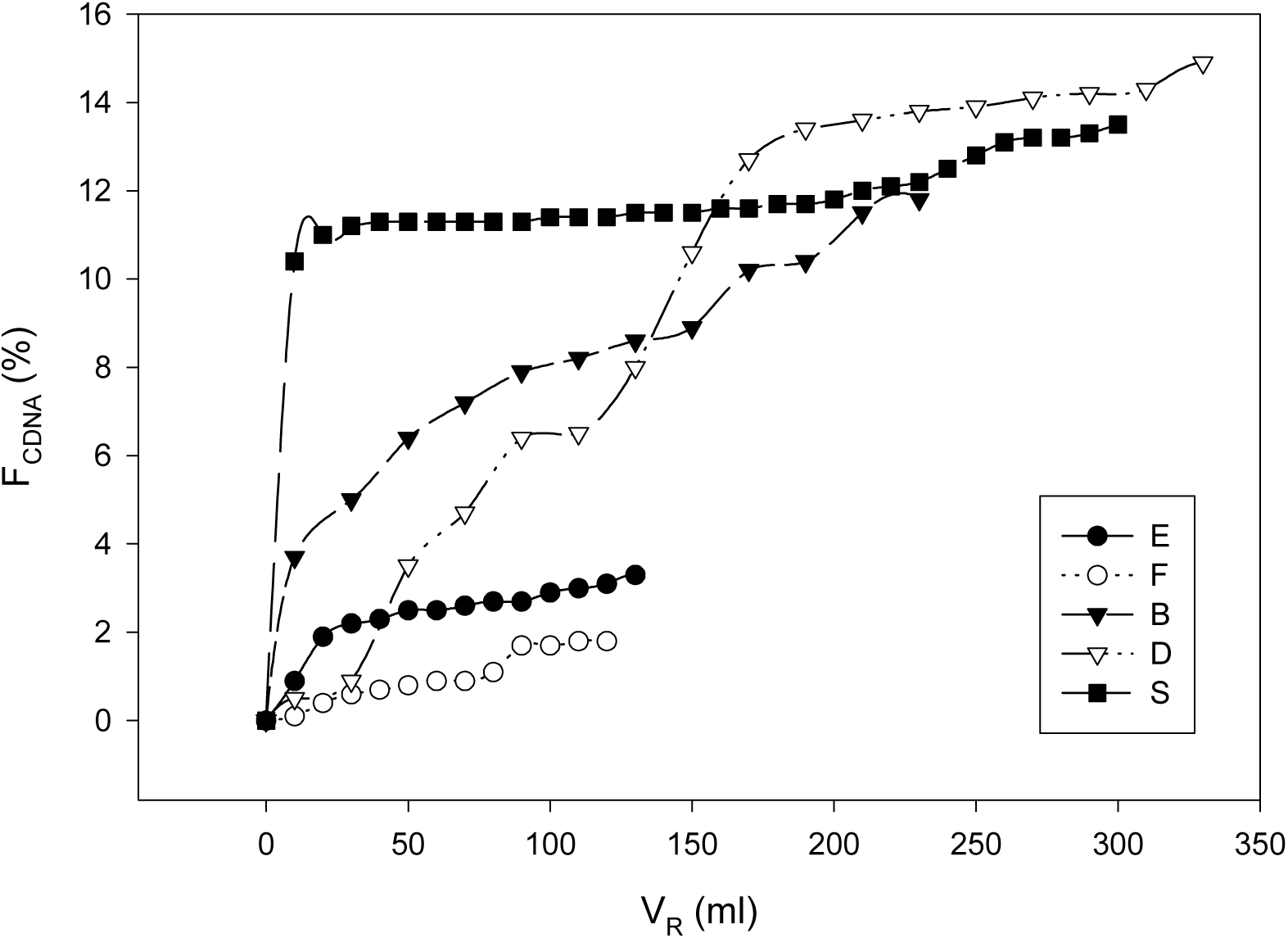
Fraction of dissolved but not absorbed felodipine (F_CDNA_ (%), normalized to 200 ml) in chyme versus recovered volume at midgut fistula, following oral administration of a micronized dose of 10 mg felodipine suspended in 200 ml of NaCl 0.9 % (#E and #F) or glucose solution 20 % (##B, D and S).

Data for bile acids were considerably lower than data reported for „hepatic bile“ in literature (Erlinger, 1987). In contrast to these studies we reported „duodeno-jejunal bile“ (and not hepatic bile) collected 76 cm distal to the pylorus after administration of 200 ml aqueous solutions. A possible “dilution effect” may explain this discrepancy. Furthermore, bile was withdrawn permanently with chyme sampling and was therefore detracted from the enterohepatic cycle (Beck, 1998).

### 3.7 Effect of fluid administered on dissolution of felodipine prior to midgut

Experimental determination of in-vivo dissolution (dissolution rate in the gut lumen) is difficult since absorption takes place simultaneously with dissolution. Nevertheless, taking into consideration very low plasma concentrations of felodipine the fraction of *absorbed* felodipine might be negligible. In the case of experiments with coarse suspensions of felodipine, concentrations of felodipine in solution at midgut were in the great majority of cases below the detection limit. The percentage of dissolved but not absorbed felodipine recovered at midgut (F_CDNA_) after coarse grade administration was never greater than 0.25 %. In general, percentage of felodipine recovered at midgut in dissolved form was higher for the micronized drug. 12 min. after administration of 200 ml saline 0.9 %, F_CDNA_ (referred to a dose of 10 mg) was determined to be 2.9 % and 1.9 %, respectively. 30 min. after administration of 200 ml glucose 20 %, F_CDNA_ was determined to be 6.4 %, 13.9 % and 12.0 %, respectively. The fluid administered appears to play a role in dissolution, with percentage dissolved recovered at midgut being substantially higher after glucose than after saline administration. It is hypothesized that this is at least partly attributable to the greater volumes generated by the administration of glucose solutions. A further possibility includes alterations in the mixing pattern. The luminal dissolution rate of felodipine was determined to be greater following coadministration with hypertonic glucose than with 0.9 % saline. This correlated well with an increased intestinal fluid volume following oral administration of 20 % glucose solution but did not correlate with an altered saturation solubility (Spearman rank sum test, P>0.100).

The correlation in figure 7 is based on the Pearson und Bravais test with R=0.972 (P<0.001) and both sets of data were normally distributed (P=0.44 and P=0.247 Kolmogorov-Smirnov). This correlation includes data from both, saline and glucose solutions, and fits well the relationship of Noyes, Whitney, Nernst and Brunner with its volume-dissolution rate linearity.

**Fig 7.**
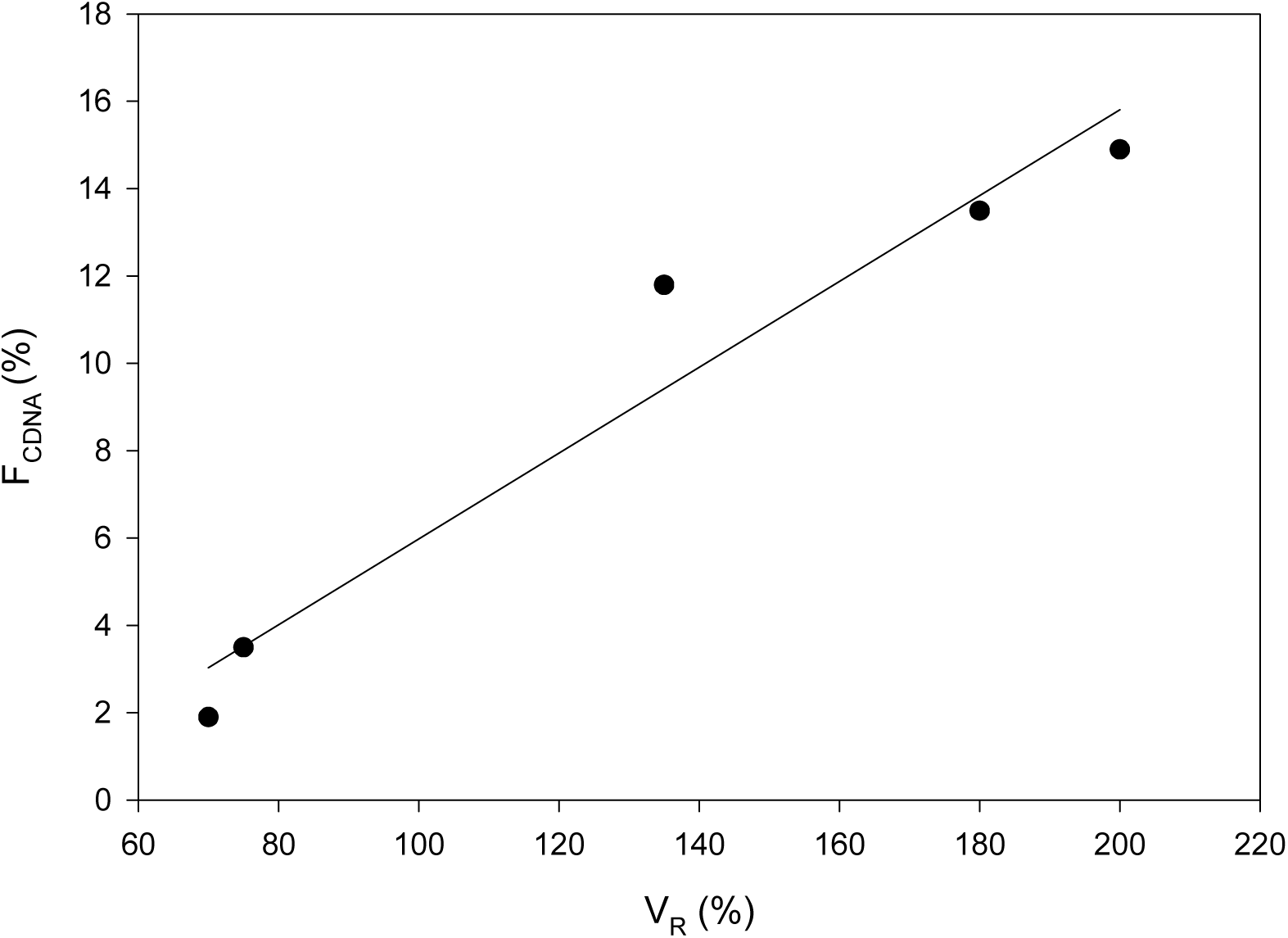
F_CDNA_(%)=0.098*(V_R_)+3.85, (R^2^=0.945, P<0.001), where F_CDNA_ (%) represents the cumulative dissolved but not absorbed fraction of felodipine in chyme, (V_R_) describes the recovered chyme fraction (volume normalized to initial 200 ml).

### 3.8 Plasma profiles

Plasma profiles are shown (pc) for Labrador Nubbe after various oral administrations (Fig. 8). Please note that these profiles are taken under regular aspiration of chyme at midgut.

**Fig. 8:**
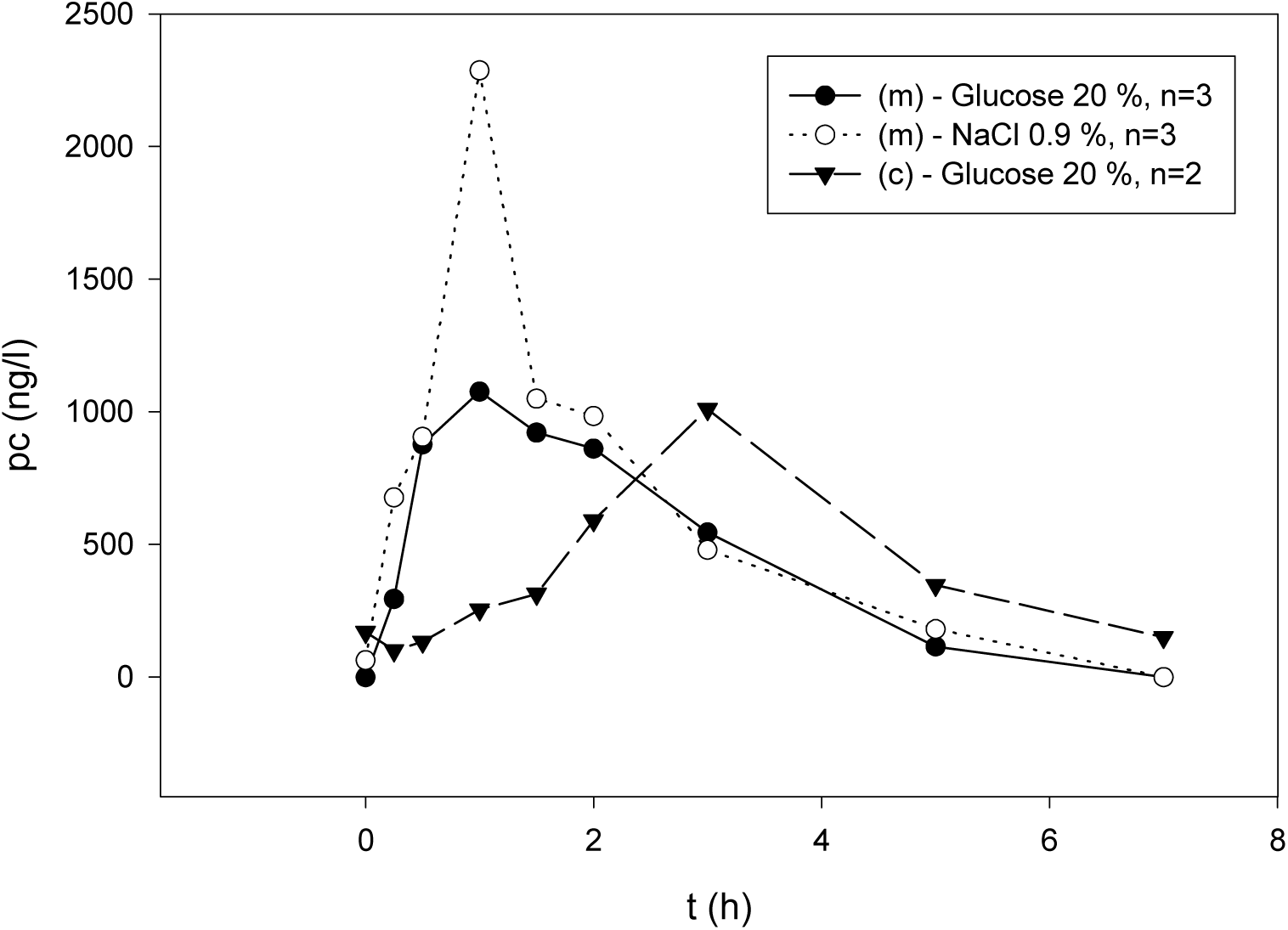
Representative mean plasma concentration (pc) profiles following the administration of 10 mg of micronized (m) or coarse grade (c) felodipine in Labradors. The drug was either suspended in 200 ml NaCl 0.9 % or glucose 20 %.

#### Normal saline versus glucose 20 %

Maximum plasma concentrations (Cmax) were higher after administration of micronized (m) felodipine suspended in normal saline (NaCl 0.9 %) compared to glucose 20 %. This is possibly an effect of a faster (“rush like”) release of non-caloric fluids from the stomach. In contrast, glucose solutions cause *a more even release* of the liquids from the dogs stomach resulting in a lower Cmax.

#### Coarse grade dose (c) versus Micronized dose (m)

Cmax of the coarse grade substance was determined to reach an equivalent magnitude compared to micronized (m) felodipine. However, in the case of glucose solution 20 %, *Tmax* was delayed considerably. This result is consistent with delayed dissolution observed in previously reported experiments using coarse grade particles (Diebold, 2000). In accordance with the law of Noyes, Whitney, Nernst and Brunner dissolution of the coarser particles progresses more slowly in vivo since the specific surface area is lower compared to micronized drug substance.

### 3.9 Plasma data

AUC_7h_ after administration of felodipine solution is 2.4 to 22.8 times higher than after administration of a suspended dose of felodipine (Tab. 1). Tmax appears earlier (0.5 to 1 h) compared to suspensions (1.33 to 3 h). Relative bioavailability (F_rel-7h_) for micronized felodipine is calculated to be 17 to 23 % and therefore somewhat higher than for the coarse grade drug (8-15 %). Absorption from suspensions is much lower than from solution. Data are highly variable for suspensions. The variability in the data may be related to the variability in lumenal solubility of felodipine (time dependency!), and may also be partly due to removal of drug at midgut via fistula. Absorption appears to depend on particle size rather than on fluid co-administered - but variability may be masking differences. The results are similar in the two dogs. In general, pharmacokinetic data are also consistent with the hypothesis that absorption continues at more distal sites (Abrahamsson, 1997).

**Tab. 1:** Plasma data for the Labradors Nubbe and Nixon following oral administration of micronized, coarse grade or dissolved felodipine (aspiration of chyme at midgut fistula). Reported plasma concentrations of felodipine are normalized on a dose of 10.0 mg of felodipine.

| Parameter |  | Nubbe<br>saline 0.9 % | glucose 20 % | Nixon<br>glucose 20 % |
| --- | --- | --- | --- | --- |
| **micronized** |  |  |  |  |
|  |  | (n=3) | (n=3) | (n=1) |
| C<sub>max</sub> | (ng/l) | 2424 | 1236 | 2759 |
| T<sub>max</sub> | (h) | 1.33 | 1.33 | 2 |
| AUC<sub>24</sub> | (ng*h/l) | 4069 | 3367 | 15483 |
| AUC<sub>7</sub> | (ng*h/l) | 4069 | 3094 | 7416 |
| F<sub>rel-7h</sub> | (%) | 18 | 17 | 23 |
| **coarse grade** |  |  |  |  |
|  |  | (n=2) | (n=1) | (n=2) |
| C<sub>max</sub> | (ng/l) | 1009 | 215 | 3641 |
| T<sub>max</sub> | (h) | 3 | 3 | 1.5 |
| AUC<sub>24</sub> | (ng*h/l) | 4450 | 1005 | 21897 |
| AUC<sub>7</sub> | (ng*h/l) | 3176 | 1397 | 4031 |
| F<sub>rel-7h</sub> | (%) | 15 | 8 | 13 |
| **solution** |  |  |  |  |
|  |  | (n=1) | (n=1) | (n=1) |
| C<sub>max</sub> | (ng/l) | 19636 | 10068 | 22478 |
| T<sub>max</sub> | (h) | 0.5 | 0.5 | 1 |
| AUC<sub>24</sub> | (ng*h/l) | 33203 | - | 36616 |
| AUC<sub>7</sub> | (ng*h/l) | 22359 | 17703 | 31880 |

## 4 Conclusion

200 ml 20 % glucose solution and 0.9 % saline solution produce dissimilar flow rates at midgut and influence gastric emptying. These fluids can be used as matrices (carriers) to alter and to access hydrodynamic effects on in vivo dissolution. The latter is also affected by various attributes of intestinal fluids (chyme). The observed time dependency of drug solubility in the gut lumen seems to play an underestimated role so far.

## ACKNOWLEDGMENT

The author would like to express his gratitude to Drs. Bertil Abrahamsson, Alfred Bayati, Jennifer B. Dressman, Edmund Kostewicz, Britta Polentarutti, Anna-Lena Ungell and Mia Wilsson-Rahmberg for their advice and assistance as well as to Jennifer B. Dressman for her kind and continuous support of the study. Support of this project by the Departments of Biopharmaceutics, DMPK and Bioanalytical Chemistry at AstraZeneca (former Astra Hässle) AB, Mölndal (Sweden) is gratefully acknowledged. The work is based on the authors book, published with Shaker Verlag, Aachen 2000 (see references).

## Notes

### Competing Interest Statement

The authors have declared no competing interest.

